# Heritable associations with microbial communities are essential for necrotrophic pathogen resistance

**DOI:** 10.1101/2020.07.16.207597

**Authors:** Cloe S. Pogoda, Stephan Reinert, Zahirul I. Talukder, Ziv Attia, Jason A. Corwin, Kennedy L. Money, Erin C.E. Collier-zans, William Underwood, Thomas J. Gulya, C. Alisha Quandt, Nolan C. Kane, Brent S. Hulke

## Abstract

Host-microbe interactions are increasingly recognized as important drivers of organismal health, growth, longevity, and community-scale ecological processes. However, less is known about how genetic variation affects hosts’ associated microbiomes and downstream phenotypes. We demonstrate that sunflower (*Helianthus annuus*) harbors substantial, heritable variation in microbial communities under field conditions. We show that microbial communities explain up to 77.5% of the heritable variation in resistance to root infection caused by the necrotrophic pathogen *Sclerotinia sclerotiorum*, and that plants grown in sterilized soil showed almost complete elimination of pathogen resistance. Association mapping revealed 69 genetic locations related to microbial abundance and *Sclerotinia* resistance. Although the genetic architecture is complex and quantitative, we have, in large part, elucidated previously unexplained genetic variation for resistance to this pathogen. This suggests new targets for plant breeding and demonstrates the potential for heritable microbial associations to play important roles in defense in natural and human-altered environments.

## Introduction

Plant-microbial interactions have long been understood to be important by ecologists ^1,2^, but the advent of agroecology and DNA methods for profiling these communities has stimulated renewed interest among scientists and plant breeders. A very important site of these interactions is the soil-root interface within the soil rhizosphere ^3,4^. Previous research has indicated that certain microbial components of the rhizosphere can be protective to the plant and enhance tolerance and resistance to biotic and abiotic stresses, benefit overall biodiversity, and increase nutrient uptake and availability ^5,6^. While research has suggested that the microbial community associated with each plant can be affected by species and even genotype,^7^ many previous reports have also suggested that differences in soil abiotic characteristics have a larger impact on rhizosphere microbial community composition than host natural genetic variation ^8–10^. Thus, a better understanding of how plant genetic diversity can shape the soil rhizosphere microbiome will affect how the microbiome is used to enhance important plant traits and soil health. Particularly intriguing are suggestions that the microbiome may play an important role in plant pathogen resistance^2^.

Cultivated sunflower (*Helianthus annuus* L.) offers an excellent system to study how plant genotype may influence plant-microbial interactions and associated disease resistance. Sunflowers are the fourth most valuable annual oil crop, with 480 million tons grown worldwide ^11^, used as a high value vegetable oil for human consumption. It is also a naturally cross-pollinated species with excellent diversity panels and associated genomic resources. One serious pathogen of sunflower, *Sclerotinia sclerotiorum*, is a necrotroph known to infect most herbaceous, dicotyledonous plants in temperate climates, including important crops such as soy, canola, lettuce, lentils, peas, and beans ^12^. *Sclerotinia*-induced basal stalk rot and wilt is the result of myceliogenic infection of the roots, which is unique to sunflowers and related species ^9^. Diseased plant symptoms include sudden wilting of the plant during seed filling and basal stalk lesions, often leading to visible signs of the pathogen including white mycelia and black sclerotized bodies, and resulting in yield loss due to plant lodging and premature death ^13,14^. Disease management in sunflower has only been successful by selective breeding for heritable resistance to the pathogen ^14,15^so a better understanding of genetics underlying resistance is essential.

Despite its clear importance, little is known about the mechanisms through which host-plant resistance is conferred. Previous studies have not identified obvious plant traits associated with resistance. With the rhizosphere being the site of infection for *Sclerotinia* basal stalk rot, we focus here on mechanisms that specifically include rhizosphere microbial community dynamics. In so doing, we identify how the non-*Sclerotinia* microbial communities affect *Sclerotinia* resistance. Together, these analyses suggest that microbial partners explain previously unexplained heritable variation in *Sclerotinia* resistance. By exploring the tripartite interaction of plant genotype, rhizosphere composition, and *Sclerotinia* disease, we provide new insight into plant defense against this cosmopolitan pathogen, which in turn will provide the potential for new methods of control that include enhancement of the host-plant genetic contributions to soil community health together with probiotic applications or improved soil microbial functioning.

## Methods

### *Helianthus annuus* Plant Material

Plant materials used in all field experiments were from the Open Pollinated Variety diversity panel of sunflower, which contains 260 USDA plant introductions (PIs), of which ten were released lines from the USDA sunflower breeding program (a list is available in ^14,16^. Of these, 95 were chosen randomly for further analysis of rhizosphere composition and another environment of inoculated basal stalk rot testing at the Carrington Research and Extension Center, Carrington, ND, USA, in 2017. Six of the released sunflower lines from the USDA sunflower breeding program were included in the subset.

For the greenhouse experiment, 40 accessions were further chosen from the subset of 95 lines (Table S1). These 40 were divided equally between highly susceptible or highly resistant to *Sclerotinia* basal stalk rot in the field.

The sunflower lines associated with each aspect of our work are listed in Table S1.

### *Helianthus annuus* DNA Extraction and Sequencing

Lyophilized leaf material for the 260 PIs were ground using tungsten carbide bearings in a Qiagen 96-well plate shaker. A total of 10 plants per PI were co-processed for each sample. Genomic DNA (gDNA) was extracted from leaf tissues using a Qiagen DNeasy 96 plant kit. The manufacturer’s protocol was modified to include the addition of 10 mM SMB (sodium metabisulfite) to the initial lysis buffer, a 45 minute incubation at 65 °C for the ground material in lysis buffer, a 100% ethanol wash before final drying of the membrane prior to elution, and DNA storage in an elution buffer that contained 10mM DTT (dithiothreitol), which have all been shown to improve DNA concentration and purity ^17,18^. Extracted samples were stored at -20 °C prior to library preparation.

Genomic libraries were prepared following standard protocols using Nextera® XT DNA library prep kits (Illumina®) and were barcoded with the Nextera® adaptors i5 and i7. Pools that passed QC were processed for an average coverage depth of 5x, 151 bp, paired-end HiSeq® reads at the NovoGene sequencing facility in Sacramento, CA.

### *Helianthus annuus* Variant Calling

Demultiplexed data was downloaded directly from NovoGene’s servers. FASTQ data were trimmed using Trimmomatic version 0.38 with the following parameters: NexteraPE-PE.fa:2:30:10 LEADING:3 TRAILING:3 MINLEN:100, with NexteraPE-PE.fa containing the standard set of Nextera adapters to be trimmed from reads. Resulting FASTQ files were aligned to the most up-to-date assembly available as of writing, HA412-HO.v2.fasta (L. Rieseberg, pers. comm., 2019). Variant calling was performed using samtools *mpileup* (samtools mpileup -I -uf) and bcftools *call* (bcftools call -v -m). A single variant call file (vcf) table was created and then pruned to contain the individuals specific to each environment/trait to remove poor quality genotypic and phenotypic data from downstream analysis.

The vcf table was filtered for single copy sites based on depth (sites with depth between 200-1000 were retained). Additionally, filtering was done to select sites with a minimum quality score of 100 (minQ=100), minor allele frequency of 0.1 (maf=0.1) and max missingness value of 0.75 (max-missing=0.75). Missing data were imputed using BEAGLE version 5.0 with default settings retained ^19^. Plink version 1.9 with the parameters: -- indep 100 5 10, corresponding to a window size of 100 SNPs, step size of 5 SNPs, and a VIF threshold of 10 (equivalent to an R^2^ value of 0.9), were used to select markers with low redundancy for use in subsequent steps ^20^.

### *Sclerotinia* Disease Incidence / Resistance Phenotyping

A previously studied, high quality, multi-environment *Sclerotinia* basal stalk rot data set was re-analyzed in our study. Details of the experiments are available in Talukder et al. (2014) ^14^. Briefly, the Open Pollinated Variety diversity panel was grown in four environments in the USA (Crookston, MN, 2008; Crookston, MN, 2009; Davenport, ND, 2008; and Grandin, ND, 2009). Field plots included about 20 plants and were arranged in a randomized complete block design with two replications per environment and 13 nested sets-in-reps incomplete blocks per replication, to effectively control error due to uncontrollable field trends ^21^. Plots were side-dressed with dried millet seed infested with *Sclerotinia* isolate NEB-274 and were rated at maturity as percentage of plants with disease symptoms in each plot. These datasets are referred to by location name and year when analyzed individually, and as the multi-environment dataset when analyzed based on multi-environment mean values.

### Rhizosphere sample collection

The analysis of rhizosphere composition was conducted in one environment (Carrington, ND) in 2017, in a field with diverse crop history and multiple years of inoculated *Sclerotinia* trials. This site was chosen because the likelihood that soil microbial populations antagonistic to *Sclerotinia* existed there was high, based on crop history and previous use as a *Sclerotinia* disease trial site. Approximately 20 plants of each PI were grown in single row, 6.1 m plots, spaced 0.76 m apart, using a Case IH 1200 series planter fitted with Almaco belt cones. The experiment consisted of a case-control design with nested randomized complete block design, such that two replications on one-half of the field area were inoculated with dried millet seed infested with *S. sclerotiorum* isolate NEB-274, and another two replications were mock-inoculated with autoclaved, dried millet only. At late bud stage, two plants from each mock-inoculated plot were destructively sampled for rhizosphere composition analysis. Soil samples were collected by extracting the root ball with a shovel (approximately 15 cm wide and 15 cm deep) and removing the loose dirt from the roots. The remaining soil adhered to the roots and was separated from the roots by shaking it into a metal pan. Roots and non-soil matter were removed, and the rhizosphere-associated soil further sized using a 2 mm aperture sieve (Seedburo). Total volume of rhizosphere soil collected per sunflower accession was approximately 50 ml. Samples were stored at -20 °C prior to DNA extraction. All plots were rated near maturity according to Talukder et al. (2014) for basal stalk rot percent incidence ^14^.

### Rhizosphere DNA and Bioinformatic Processing

DNA from rhizosphere-associated soil was extracted using the standard protocol for the Qiagen DNeasy PowerSoil HTP 96 kit. The phylogenetically informative prokaryotic 16S were amplified using the barcoded 515 f/926r and ITS1F/ITS2 primer pairs, respectively. Sample amplicons were pooled and sequenced on an Illumina MiSeq instrument using paired end sequencing for 251 bp at the BioFrontiers sequencing center at the University of Colorado, Boulder.

Raw reads were processed using a previously described uSearch bioinformatic pipeline ^22^. Briefly, primer dimers and adapter sequences were removed using the ‘cutadapt’ software and filtered for high quality reads using USEARCH v10 ‘fastq_filter’ function ^22,23^. Paired-end reads were merged using USEARCH v10 ‘fastq_mergepairs’ function and samples were demultiplexed using a custom in-laboratory python script. The sequences were de-replicated and denoised using the ‘derep_fulllength’ and ‘unoise3’ algorithms, respectively. Raw reads were mapped back to the unique denoised sequence database and an operational taxonomic unit (OTU) count table was created using custom python scripts. Putative phylogeny of unique denoised sequences was determined using the RDP classifier against the Greengenes phylogenetic database v13.5; for 16S sequences ^24^. Any sequences that mapped to eukaryotic chloroplast or mitochondrial sequences were removed from the analysis.

### Greenhouse validation experiment

Native soil was obtained in late 2018 from the same field as the original rhizosphere study near Carrington, ND. The soil was placed in plastic boxes and transported to the USDA-ARS in Fargo, ND. The soil inside the boxes was broken up with a shovel and large plant debris and rocks were removed. Half of the soil was marked for autoclaving, with the remaining half not autoclaved. Autoclaved soil was treated twice for 45 minutes on a wet cycle at 121 *°*C, with agitation between cycles. To measure the effectiveness of the autoclave sterilization, *Geobacillus stearothermophilus* bioindicators (Crosstex, Little Falls, NJ) were placed in the middle of the bags of soil. The vials were removed during agitation and placed back in the center of the soil for the second round of autoclaving. After the two cycles of autoclaving were complete, twenty bioindicators and two control indicators were activated and incubated for 24 hours at 60 *°*C, as prescribed by the manufacturer. The treatment bioindicators were compared to the controls and the results showed that two of the twenty were not sterilized. The soil then underwent a third 121 *°*C wet cycle for 30 minutes, with five bioindicators placed randomly in the soil. These vials were evaluated after the 24 h incubation period and indicated that the soil had been effectively sterilized.

The experimental design was based on the procedure of Underwood et al. (2020) ^25^. All plants were grown from seed in the greenhouse. Initially, 36 plants of each of the 40 accessions were planted in each of the native and the autoclaved soil, in cell flat trays (TO Plastics 730654C, Clearwater, MN, USA). Plants grown in autoclaved soil and native soil were physically separated on different benches. These plants were allowed to grow approximately 5 weeks after planting with up to twice-a-day watering to avoid water stress.

After 5 weeks, the plants were removed from the early growth trays. They were transplanted into new trays in a split-plot design with three replications nested within the main plot treatment of autoclaved vs. native soil, to prevent crossover inoculation across soil treatments within a replication. Each replication was completely randomized, containing 12 plants each of the 40 accessions, and placed in fifteen 32-cell trays with 2.5 mL (or about ∼0.38 g) of *S. sclerotiorum* inoculated millet (isolate NEB-274) at the bottom of each plant. Development of millet inoculum has been previously described ^14^. Each replicate was placed on a separate bench in the same greenhouse.

After inoculation and randomization, for the next 28 days, each plant was evaluated daily for symptoms of *Sclerotinia* basal stalk rot. Plants were evaluated for days-to-death by noting meristematic necrosis, typified by browning of the apical meristem / emerging leaves, and in most cases, permanent terminal wilt. If a plant exhibited either or both of these symptoms for two consecutive days, it was noted as dead on the first day and removed from the tray the second day.

### Statistical Analysis

#### Disease incidence / resistance

Percent incidence data for all basal stalk rot experiments were analyzed using PROC MIXED of SAS v. 9.4 (SAS Institute, 2013), with all effects random except environment, which remained fixed^26^. Phenotypes were inverted in some analyses for scaling purposes, and the inverted phenotypes were called percent resistance. Variance components were used to compute broad-sense heritability on an entry-mean basis, using the formula H^2^ = V_g_/(V_g_ + V_ge_/l + Vε/rl) for multiple environments and H^2^ = V_g_/(V_g_ + Vε/r) for single environment analyses, where V_g_, V_ge_, and Vε are the accession, accession-by-environment, and error variances, respectively; r is the number of replications per environment; and l is the number of environments^27^.

#### OTU abundances

We filtered our dataset for all samples with less than 5,000 OTU counts and OTUs that occurred in less than 25% of samples were removed, as their inclusion could skew downstream statistical analysis. The resulting filtered OTU table was normalized using the trimmed M-means algorithm using the ‘edgeR’ package within the R statistical environment ^28,29^. PERMANOVAs of the normalized pseudo-count matrix were obtained from adonis() in the ‘vegan’ package of R using the following model: Y_rg_ = R_r_ + G_g_ + ε_rg_, where Y_rg_ is the normalized pseudo-count, R_r_ is the field replicate, G_g_ is the sunflower accession, and ε_rg_ is the error effect ^30^. The effect of plant accession on each individual OTU was also obtained with a negative binomial generalized linear model using the same model for each individual OTU and the resulting p-values were FDR corrected for multiple comparisons. Broad-sense heritability across the sunflower accessions for each OTU was defined as H^2^ = V_g_/(V_g_ + Vε/r). Model-corrected means for each OTU and sunflower accession were determined from the nb.glm using the LSmeans function within the ‘doBy’ package.

#### Network analysis and association with disease

Bacterial OTUs that may serve as a protective function against *Sclerotinia*-induced stalk rot were first extracted using network analysis. Briefly, model-corrected means for each sunflower accession and OTU were correlated with the model-corrected means of basal stalk rot from the Carrington 2017 inoculated study. OTUs were chosen that were most negatively correlated with incidence of stalk rot with an FDR corrected p-value < 0.0001. This analysis resulted in a total of 42 bacterial taxa negatively associated with incidence of *Sclerotinia* infection. NMDS plots were generated using R-package *vegan* in R version 3.6.

Each of the 42 bacterial taxa that were negatively correlated with *Sclerotinia* incidence were then tested individually against resistance phenotypic data using a general linear model (‘anova’ function in the ‘lm’ package of R) ^29^. The model p-values were corrected using the Bonferroni method, and OTUs significant at the corrected-p < 0.05 threshold were tested for pairwise interaction in a series of models that consisted of all individually significant OTUs as main effects plus one interaction effect of each pair of significant OTUs. Resulting p-values were again Bonferroni-corrected by the number of interactions tested. R^2^_P_ values were extracted from the regressions.

The percentage of the heritable portion of the disease resistance phenotype explained by the heritable portion of the contribution of rhizosphere was calculated as R^2^_G_ = R^2^_P_ H^2^_SR_ H^2^_OTU_, where: R^2^_G_ is the percentage of the heritable portion of the stalk rot disease phenotypic variation explained by the heritable portion of variation in a single OTU, R^2^_P_ is the percentage of the phenotypic variation in stalk rot disease in a given environment explained by the phenotypic variation in each OTU, as above, H^2^_SR_ is the heritability of the stalk rot disease phenotype in a given environment or multi-environment mean, as calculated above, and H^2^_OTU_ is the heritability of the OTU phenotype, as calculated above.

#### Greenhouse validation experiment

Analysis of the greenhouse experiment was conducted using PROC MIXED of SAS v. 9.4 ^26^. Data were modelled as follows: Y_ijkl_ = S_i_ + R[S]_ij_ + G_k_ + SG_ik_ + ε_ijk_ + w_ijkl_, where Y_ijkl_ is the number of days to death, post-inoculation, S_i_ is the effect of i^th^ soil type (native vs. autoclaved), R[S]_ij_ is the effect of the j^th^ replication (1, 2, 3) nested within the i^th^ soil type, G_k_ is the effect of the k^th^ sunflower accession (1 to 40), SG_ik_ is the interaction of the i^th^ soil type with the k^th^ sunflower accession, ε_ijk_ is the residual model error, and w_ijkl_ is the within treatment effect of the l^th^ plant (1 to 12) nested within the i^th^ soil type, j^th^ replication, and k^th^ sunflower accession.

Least square means were calculated for each level of the soil-by-sunflower-accession interaction, using the LSMEANS option. Tukey’s multiple comparison method was used to compare the least square means. Least square means of days to death for each accession, separately for native and autoclaved soil, were then tested individually against the 42 OTUs using a general linear model (‘anova’ function in the ‘lm’ package of R), in the same way as the field data ^29^. The percentage of the heritable portion of the days-to-death phenotype explained by the heritable portion of the contribution of rhizosphere was also calculated in the same manner as the field stalk rot evaluations.

#### Association mapping

Sunflower quantitative trait nucleotides (QTN) associated with each OTU and disease phenotype were identified using the R-package *GAPIT* and specifically the *FarmCPU* function^31^. Least square means of abundance for each OTU, disease incidence of SR at each field environment (including greenhouse experiments) and for the multi-environment mean were the dependent variables used in the analyses. The first principle component, with a maximum number of iterations equal to 10 was used to detect associations between effective markers and traits. Extreme outlier genotypes were culled from the dataset based on visual observation of the principle component graph (Supplemental Figure S5). In order to identify QTNs of interest, we calculated the q-value (estimation of false discovery rate) using the R-package *qvalue* and selected sites above a threshold q-value = 0.5^32^. These QTNs were marked as potentially interesting drivers of their respective traits and environments.

#### Linkage Disequilibrium

To understand the extent of linkage disequilibrium (LD) in the vicinity of significant markers, we utilized two separate methods. First local LD was assessed chromosome-wide (for all 17 chromosomes) to identify the extent of the LD blocks due mainly to physical linkage and other population genetic forces for each of the significant QTNs. For that, we pruned the variant call file (vcf; genotypic data) of each chromosome to include every hundredth marker as well as the QTNs of interest (described above in the association mapping section) associated with each chromosome. The resulting pruned vcf table was converted into plink formatted files using vcftools. Next, plink version 1.9 was utilized to determine the R^2^ LD matrix ^20^. This matrix was further filtered to only include the pairwise comparisons for each of the significant QTNs to all other markers on the same chromosome. The resulting chromosome-wide data were graphed using *ggplot2* in R version 3.6.0 ^29^. These data were parsed to determine the size of the LD block surrounding each QTN (Table S2). Second, to find longer range LD among significant QTN within and among chromosomes, due to selection forces on unlinked loci, we used Haploview Desktop version 4.2 with default settings to examine LD among the significant QTNs of all field sites and significant OTUs ^33^.

## Results

### Identification of Rhizosphere Taxa-of-interest and Associated Heritability Values

16S marker profiling of the sunflower rhizosphere microbiome identified 890 unique bacterial and archaeal OTUs in the collected sunflower samples. Of these 890, statistical microbial diversity analysis revealed 42 taxa which are negatively correlated with disease incidence but are relatively unstructured among the 890 OTUs identified (Figure 1) and are from a variety of different phyla (Table S3). Due to their putative importance to inhibiting *Sclerotinia* establishment on the host, we focused on these 42 OTUs for further study.

**Figure 1:**
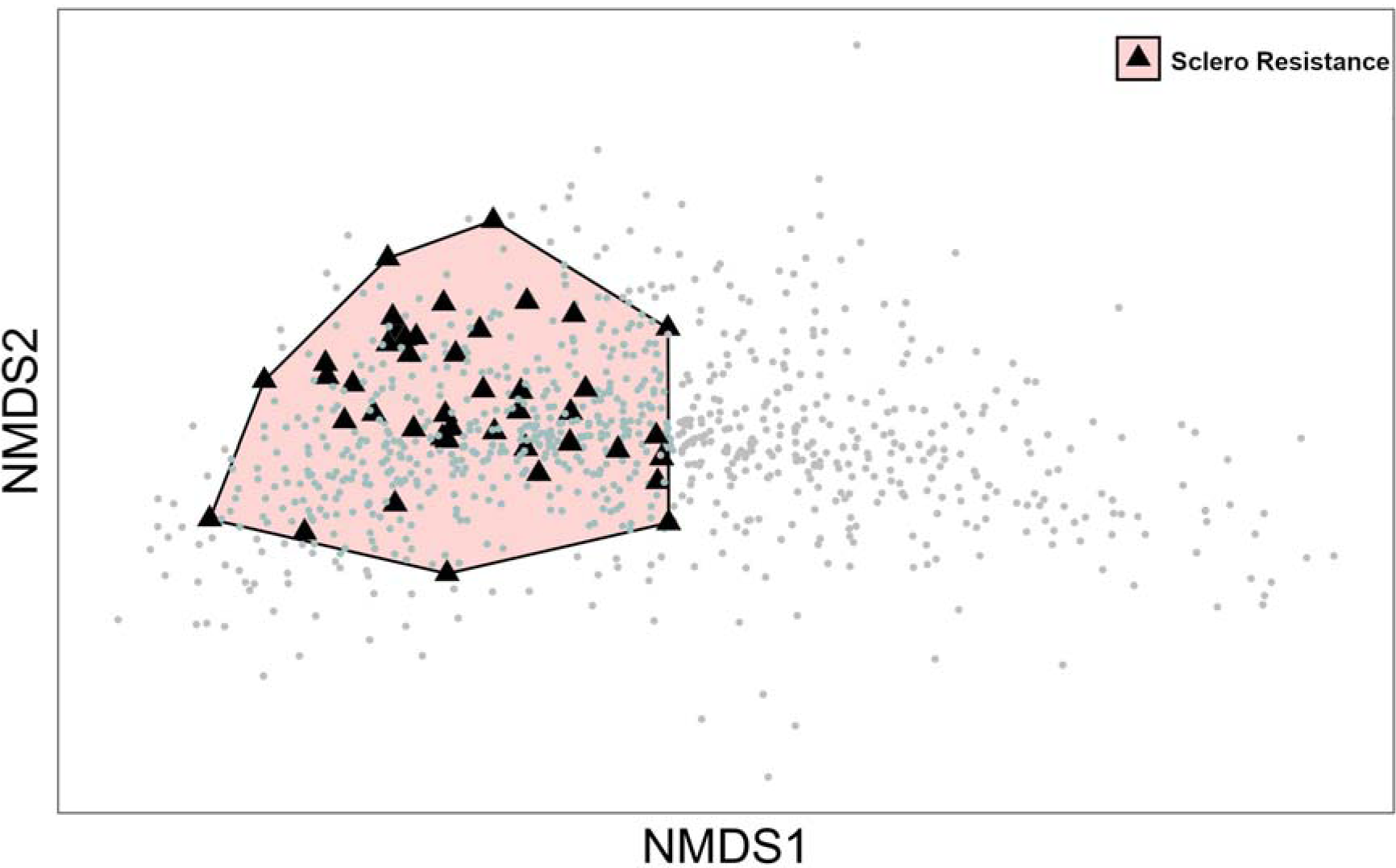
NMDS plot of the 890 (grey dots) operational taxonomic units identified in the soil sampled from Carrington, ND, in 2017. Forty-two taxa (black triangles) that are negatively correlated with *Sclerotinia* basal stalk rot disease resistance are highlighted in black. Black polygon indicates the boundaries of the 42 taxa.

Individual regression analysis of the 42 OTUs associated with disease resistance identified four bacterial taxa which showed a statistically significant effect on basal stalk rot resistance in Carrington (adjacent, *Sclerotinia*-inoculated field plots in the same environment as the rhizosphere assessment; Figure S1, Table S4). These OTUs were identified as OTU77: Acidobacteria, OTU293: Verrucomicrobia bacteria, OTU750: Rhizomicrobium bacteria, and OTU908: Actinobacteria. When OTU abundance in Carrington 2017 was regressed on the multi-environment data set of Talukder et al., 2014 ^14^, OTU750 showed a highly significant response (Bonferronized p = 0.003). When the multi-environment data from the Talukder study was reduced to individual environments, OTU293, OTU750, and OTU908 were significant in at least two of the four environments. This result shows the conservation of these relationships over a 250 km distance among several field sites and with different crop history. The broad sense heritability values for the four significant OTUs were centrally distributed among the values for all 890 individuals initially identified (Figure 2).

**Figure 2:**
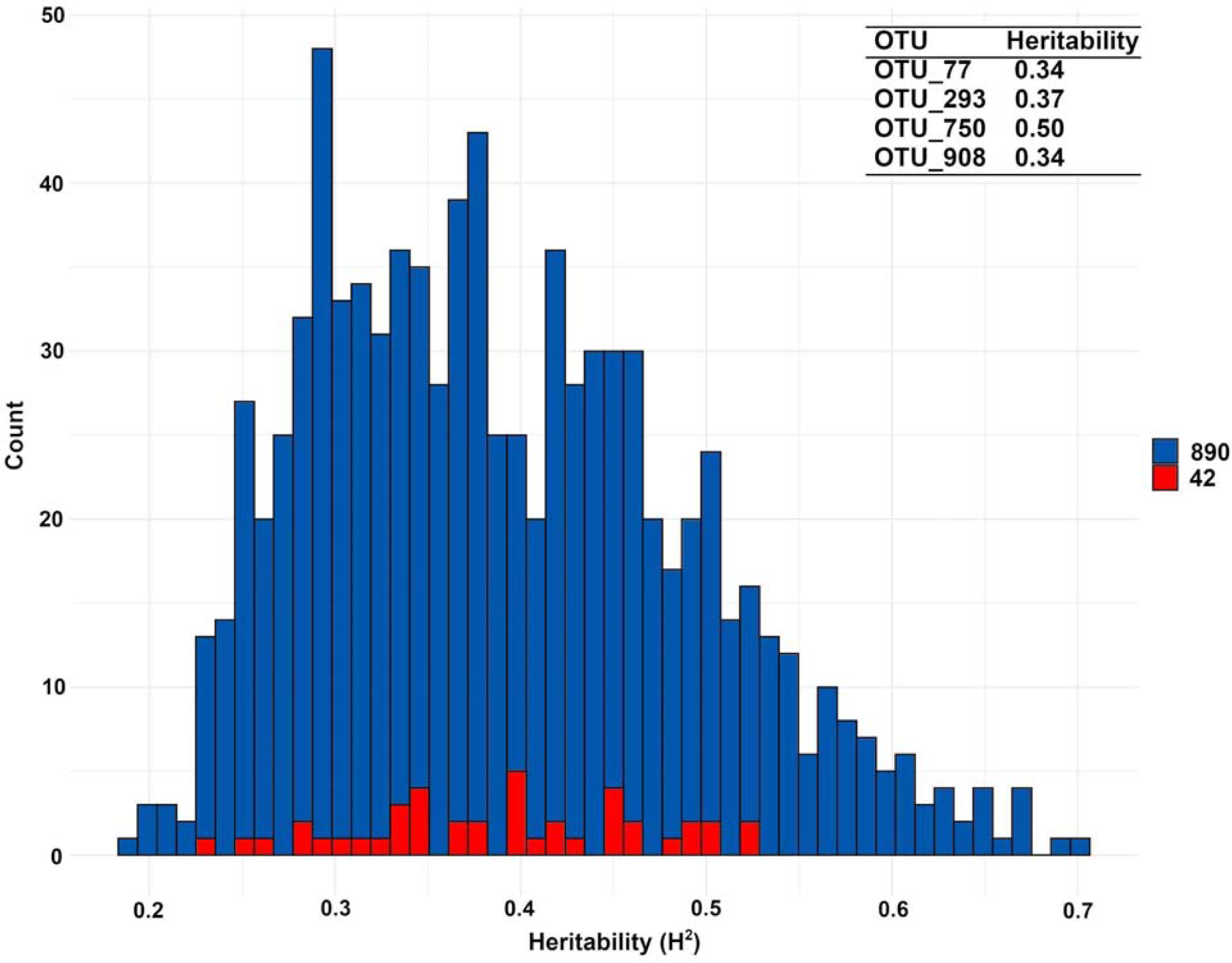
Heritability plot of the 890 bacterial operational taxonomic units sampled in 2017 from the Carrington, ND, field. Forty-two taxa that were negatively correlated with *Sclerotinia* disease resistance fall within the bars highlighted in red. The heritability values of the four statistically significant OTUs are given in the figure legend (top right).

### Variation for Basal Stalk Rot Resistance Explained by Rhizosphere

In order to better understand the role of genetics in the stalk rot phenotype, we calculated the broad-sense heritability of disease resistance for each environment studied. The broad-sense heritability estimate for *Sclerotinia* resistance in the Carrington 2017 field environment was high (0.76), while for the multi-environment mean of the Talukder et al., 2014 study, the estimate was 0.70 ^14^. Individually, those four environments were as follows: Crookston in 2008, 0.57; Crookston in 2009, 0.47; Davenport in 2008, 0.39; and Grandin in 2009, 0.35.

We then correlated the rhizosphere community present at the Carrington 2017 environment with the disease resistance phenotypes in all of our evaluations (Figure 3 & S2). We note that the only portion of phenotypic variation that could co-vary across environments was the heritable portion. Table 1 shows the percentage of the heritable portion of the phenotypic variance in stalk rot resistance explained by the heritable variance for each OTU. The percent heritable variation that was explained by each of the four OTUs in each separate field environment ranged between 11.0% for OTU77 in Grandin in 2009 and 77.5% for OTU293 in Davenport in 2008.

**Table 1:** Percent heritable *Sclerotinia* basal stalk rot resistance variation explained by each operational taxonomic unit in field and greenhouse environments.

| Taxon | Individual evaluation sites |  |  |  |  | Carrington <sup>a</sup> |  |  |
| --- | --- | --- | --- | --- | --- | --- | --- | --- |
|  | Crookston08 | Crookston09 | Davenport08 | Grandin09 | Multi <sup>b</sup> | Field | Autoclaved | Native |
| | $R^2_G$ | | | | | | | |
| OTU77 | 40.9 | 14.2 | 42.2 | 11.0 | 31.1 | 61.1 | 10.6 | 12.9 |
| OTU293 | 67.7 | 17.6 | 77.5 | 23.5 | 28.9 | 37.9 | 57.5 | 33.3 |
| OTU750 | 57.2 | 17.8 | 66.5 | 28.1 | 45.7 | 38.7 | 33.6 | 37.1 |
| OTU908 | 73.3 | 66.5 | 41.7 | 16.6 | 31.8 | 58.5 | 67.0 | 56.4 |

<sup>a</sup> Field: *Sclerotinia* field trial adjacent to rhizosphere collection experiment; Autoclaved: Greenhouse study using autoclaved Carrington soil; Native: Greenhouse study using non-autoclaved Carrington soil.
<sup>b</sup> Multi: Mean of *Sclerotinia* basal stalk rot disease incidence across four field sites (Crookston08, Crookston09, Davenport08, Grandin09).

**Figure 3:**
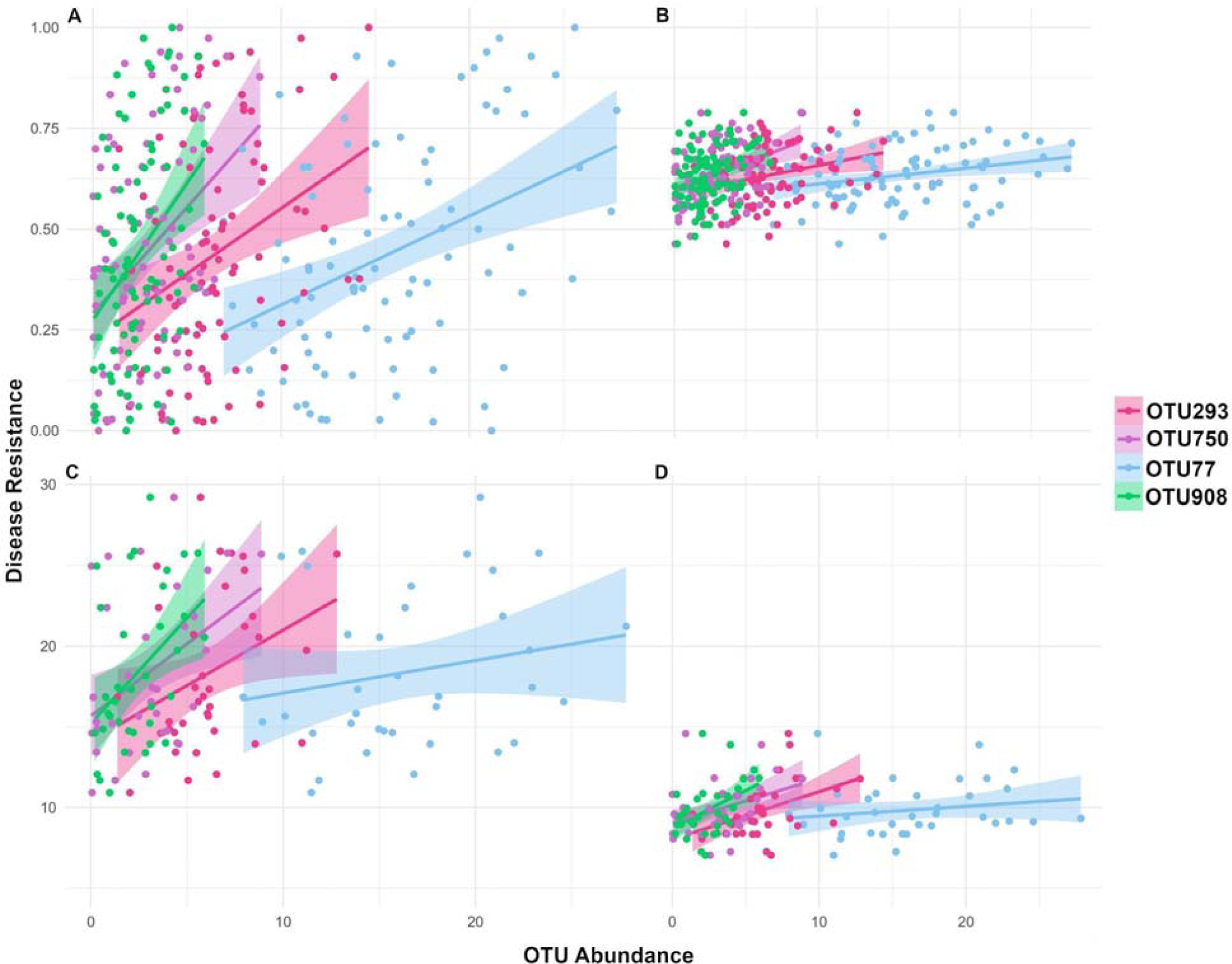
Correlation between disease resistance and operational taxonomic unit (OTU) abundance for the top four OTUs that were significant in multiple field environments. Data points are colored to indicate the OTU they represent as indicated by the figure legend. The x-axis is relative OTU abundance (count in thousands) in Carrington, 2017, for each panel. Y-axis is disease resistance calculated as 1 minus [disease incidence] (disease incidence: plants showing wilt or lesions) in panels A and B and days-to-death in panels C and D. Regression bands indicate a 95% confidence interval. **A)** Carrington field environment **B)** Multi-environment mean (combined data from four separate sites: Crookston08, Crookston09, Davenport08 and Grandin09) **C)** Greenhouse native soil **D)** Greenhouse autoclaved soil

To validate these findings, we used a greenhouse test for basal stalk rot resistance screening that has been previously verified to replicate field resistance ^25^. Forty sunflower accessions from the 95 that were sampled for rhizosphere composition in Carrington were re-evaluated in replicated greenhouse trials under both native soil and autoclaved soil conditions, with high correspondence between the field resistance and greenhouse native soil experiments (Figure 3; Table S5). Results showed that days to death from stalk rot under native soil for each sunflower accession was between 2 to 19 days longer for native soil compared to autoclaved (mean = 9 d, median = 8 d, p < 0.0001). The distribution in days to death in autoclaved soil was considerably narrower than under native soil, which resulted in a significant soil-by-accession effect (p < 0.0001). The most resistant accessions in the Carrington 2017 test survived, on average, 10 d longer in native soil than autoclaved (median = 10 d) in the greenhouse test, while the least resistant from the field test averaged 7 d (median = 6 d) longer. Despite being autoclaved, a small but significant effect of OTU on resistance occurred in that treatment, which could be due to environmental contaminants that increased over the 5 week plant growth interval before inoculation or increase from some very low level of microbial inoculum that survived autoclaving.

### Genetic architecture of the rhizosphere - *Sclerotinia* resistance association

We examined the *Sclerotinia* and rhizosphere datasets for significant QTNs using standard GWAS techniques detailed above. Due to moderate heritabilities of both traits and quantitative inheritance, multiple genomic loci of variable effect size were associated with each OTU and disease screening environment. Markers that fell above a q-value threshold of 0.5 were identified for further analysis, which resulted in 69 QTNs total for all traits and environments (Figure 4, 5, S3). For each QTN, local LD tended to decay dramatically in most cases. There were notable exceptions, including chromosomes 3, 6, 7, 8, 9, 10, 13, 14, 16 and 17, which had LD at significant QTN extending beyond 1 Mbp (Table S2). Chromosome 9 had the most QTNs at 7, while chromosome 5 had the fewest at 1 QTN.

**Figure 4:**
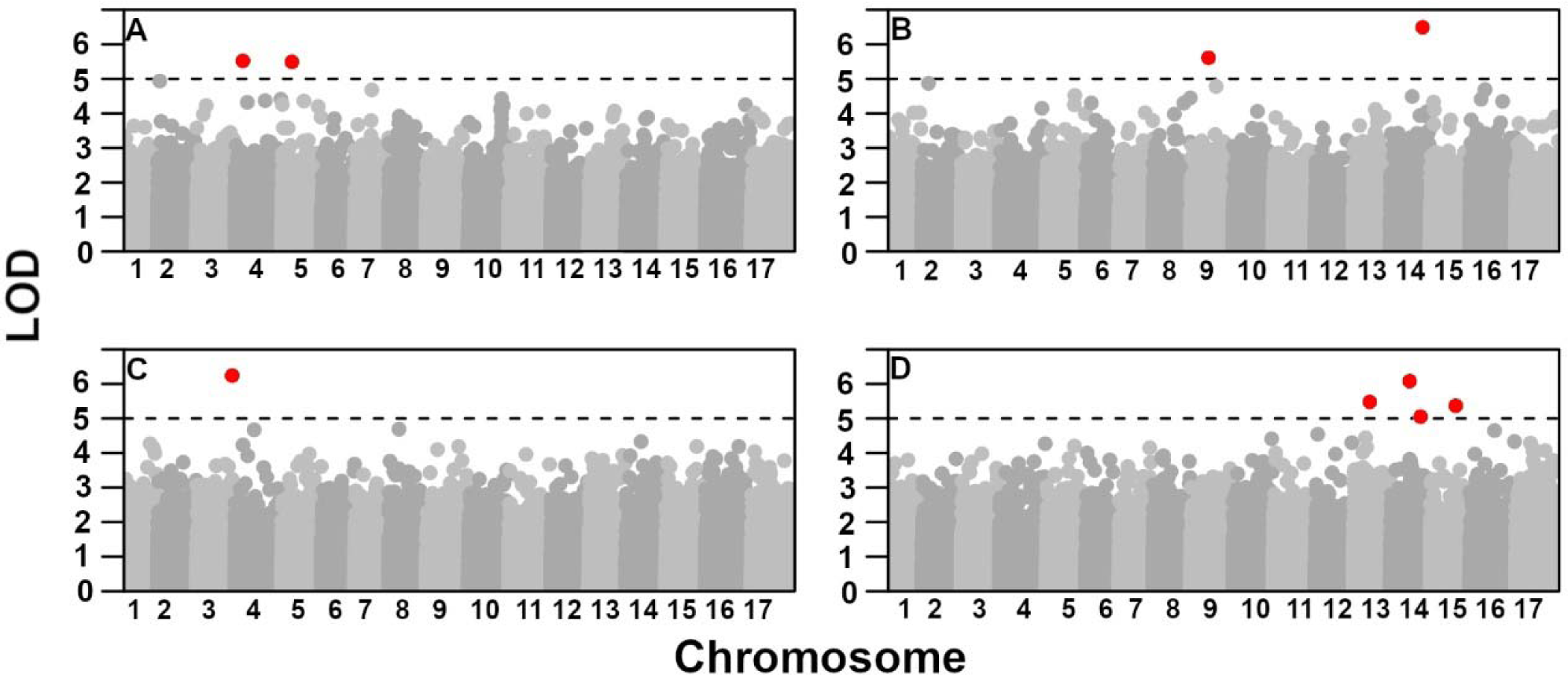
Genome-wide Manhattan plot for the abundance of four operational taxonomic units (OTUs) that are significant in multiple environments. Chromosomes are labeled and represented in alternating greyscale. The marker sites above a q-value of 0.5 are highlighted in red for each plot. **A)** OTU77 **B)** OTU293 **C)** OTU750 **D)** OTU908

**Figure 5:**
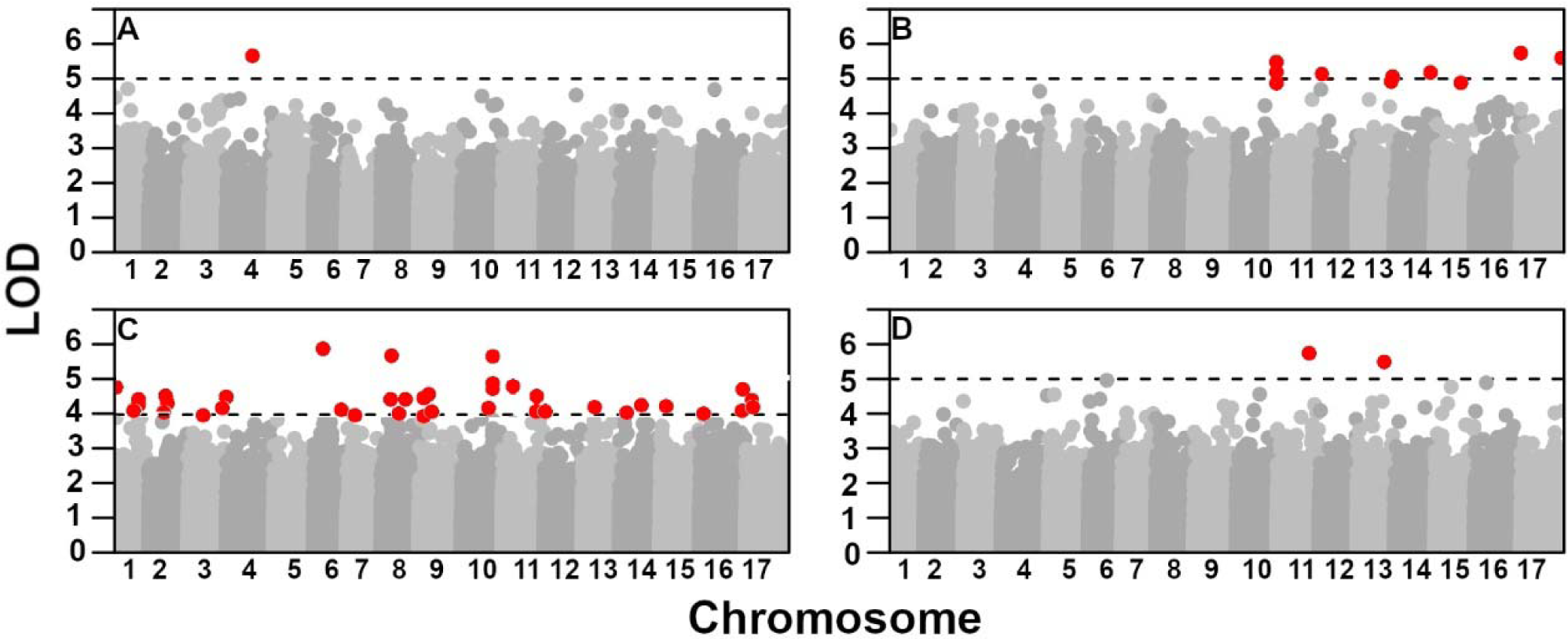
Genome-wide Manhattan plot for stalk rot resistance in four experiments. Chromosomes are labeled and represented in alternating greyscale. The marker sites above a q-value of 0.5 are highlighted in red for each plot. **A)** Carrington 2017 **B)** Multi-environment (mean of Crookston08, Crookston09, Davenport08 and Grandin09) **C)** Greenhouse native soil **D)** Greenhouse autoclaved soil

In order to identify marker-trait QTNs in high LD due to historical selection forces on the diversity panel, we calculated LD for all significant sites and focused solely on those sites which were in LD at R^2^ ≥ 0.6. Even after previous culling of collinear QTN due to local LD, there was still evidence of extensive interchromosomal LD, including 20 instances of QTN for both *Sclerotinia* and OTU phenotypes in high LD (Table S2, Figure S4). Since many of the open pollinated varieties on the panel diverged ca. 500 years ago with the diaspora of sunflower from North America overseas, it is likely that a bottleneck, followed by a population expansion, has influenced our diversity panel at loci of functional importance under cultivation or natural environments. Logically, this would include *Sclerotinia* resistance, as the disease occurs worldwide. In this scenario functional loci within each haplotype may have been maintained in high LD while neutral, intervening nucleotides underwent mutation and drift. Additionally, high interchromosomal LD was also observed among many pairs of QTN, which is almost certainly due to selection, given negligible population structure in the panel (Figure S5). Given the complexity of this LD, we subjected the LD between QTN to a neighbor-joining tree decomposition to determine networks of loci working in tandem for our phenotypes of interest (Figure S6). It is apparent that multiple, physically unlinked loci have been selected in tandem across the genome for the purpose of increasing abundance of our four OTUs of interest (and likely others), reducing *Sclerotinia* stalk rot disease (a known target of sunflower breeding since domestication), and perhaps other, yet unknown purposes.

## Discussion

Using a population of cultivated sunflower genotypes that were grown in multiple environments and evaluated for resistance to *Sclerotinia sclerotiorum* myceliogenic root infection, also known as basal stalk rot, we show a strong link between host-plant resistance and heritable associations with the rhizosphere microbial community. We hypothesize that specific bacterial OTUs increase resistance to this plant pathogen. It has been shown that healthy rhizosphere communities can contribute to not only overall plant vigor but ability to resist biotic stressors ^35–37^. By connecting the disease resistance of specific sunflower genotypes with bacterial strains present in a healthy rhizosphere, we achieve greater understanding in how to fight this destructive and detrimental pathogen.

In our study, we sampled soil from one field site (Carrington) which had a rich history of *Sclerotinia* disease studies on several different crops. We identified 890 distinct bacterial OTUs in the soil sampled. Among these 890 individuals, we further identified 42 OTUs that were significantly and negatively associated with *Sclerotinia sclerotium* disease incidence. Regression analyses identified four of these 42, individually, as being statistically significant associations. While we did not directly measure the soil communities in the other field environments we analyzed (Crookston08, Crookston09, Devenport08, Grandin09, autoclaved greenhouse soil, native greenhouse soil), we observed the same pattern of correlation in all of the field environments for some or all of these four OTUs (Figure 3 & S2), suggesting that plant genetics associated with microbial abundance and disease resistance in Carrington was also associated with disease resistance in these other environments.

The heritability values of the four OTUs identified were centrally distributed, indicating that these bacteria are truly significant and not spuriously correlated due to statistical bias. We identified the four most significant OTUs as bacteria that are commonly present in healthy rhizospheres from a wide range of environmental conditions ^38^. The ubiquity of these species in healthy plants’ microbiomes suggests that these taxa may be of general importance and not specific to the confines of our study, though further validation will be necessary to support this. Nevertheless, it supports the hypothesis of a healthy bacterial community increasing resistance to biotic stress in the common sunflower ^38–44^.

Our field results were validated with a greenhouse experiment where we grew a subset of the plant genotypes in both autoclaved (sterilized) soil and unsterilized soil directly from the Carrington field. Greenhouse experiments in unsterilized (native) field soil showed similar results to field conditions, but in sterilized soil resistant plant genotypes performed much worse in the *Sclerotinia* assay, to the extent that nearly all resistance was eliminated, thus adding to the data that a healthy rhizosphere plays an important role in a plant’s ability to resist infection ^45–47^.

Quantitative trait nucleotides (QTN) in the host plant contributing to OTU abundance and *Sclerotinia* disease incidence formed LD clusters inter-chromosomally. This provides a solid basis to assert that some of the same genetic loci that affect OTU abundance in sunflower are also resulting in disease resistance. This could be either by a pleiotropic effect (the underlying genes provide a product that enhances certain OTUs but is detrimental to *Sclerotinia*) or because of a causative effect of genes enhancing OTUs and the OTUs, in turn, inhibiting *Sclerotinia*. Our greenhouse work, already discussed, suggests the latter is more likely. Additionally, LD blocks identified between significant QTNs on chromosomes 3, 4, 9, 11, 14 and 17 are supported by previous studies which found *Sclerotinia* resistance loci on these chromosomes ^48–50^. Given the diversity panel is composed of cultivated sunflower lines at various levels of improvement and different geographic origins, high LD among QTN is likely the result of artificial and natural selection for OTU abundance together with *Sclerotinia* disease resistance. This is bolstered by the negligible signals of population structure (Figure S5). While we reason that pleiotropy is occurring, epistasis and local clustering of genes with important additive effects into haplotypes could also be important.

## Conclusion

The microbial taxa analyzed here hold promise as probiotics that could reduce the risk of *Sclerotinia* disease or as indicator organisms for improvement of agricultural soil health. More broadly, our demonstration that plant genotype shapes abundance of numerous taxa in the soil microbial community indicates a high potential for selection to act on this genetic variation, enabling evolution of more favorable plant-microbe interactions. If these bacterial associations provide benefits or costs to plant fitness, by provisioning disease resistance or other phenotypes, natural and artificial selection would be expected to shape microbial associations, either in a plant breeding context or in natural ecosystems. The heritable microbiome variation in this cultivated sunflower diversity panel suggests the potential for tradeoffs that are leveraged differently in each selection environment, but this panel also provides a valuable system with which to investigate the ecology, genetics and physiology of microbial interactions, with numerous applications as well as importance for basic science.

## Supporting information

Supplemental Figures with captions

## Acknowledgements

The authors would like to thank the staff of CHS Sunflower, Croplan Genetics, and the NDSU Carrington Research and Extension Center for providing space for our field experiments, with special thanks to the Carrington Research and Extension Center for soil for the greenhouse experiment. The assistance of multiple students and technicians in conducting this work is also acknowledged, particularly Christopher Misar, Brady Koehler, and Michael Grove. Noah Fierer and Jessica Henley provided important advice and assistance. This work was supported by USDA-ARS CRIS projects 5442-21220-024, 3060-21000-039, and 3060-21000-043; and the USDA-ARS National *Sclerotinia* Initiative. This research was also supported by BARD, the United States - Israel Binational Agricultural Research and Development Fund, Vaadia-BARD Postdoctoral Fellowship Award No.FI-577-2018.

## Author Contribution Statement

BSH, NCK, and TJG designed the study and obtained funding; ZIT, WU, and TJG managed field studies and collected *Sclerotinia* field resistance data; SR and BSH collected rhizosphere samples; KLM and BSH managed greenhouse studies and collected greenhouse data; ZIT and BSH analyzed field and greenhouse sunflower disease data; JAC, CSP, and CAQ analyzed rhizosphere samples and associated data; ECEC and CSP sequenced sunflower lines and developed the genomic marker dataset; CSP and SR conducted genome-wide association studies; CSP, SR, ZA, BSH, NCK, and CAQ initially drafted the manuscript; and the manuscript was revised and all aspects approved by all authors.

