## Supplementary figures and images for "Heritable associations with microbial communities are essential for necrotrophic pathogen resistance"

### Figure S1

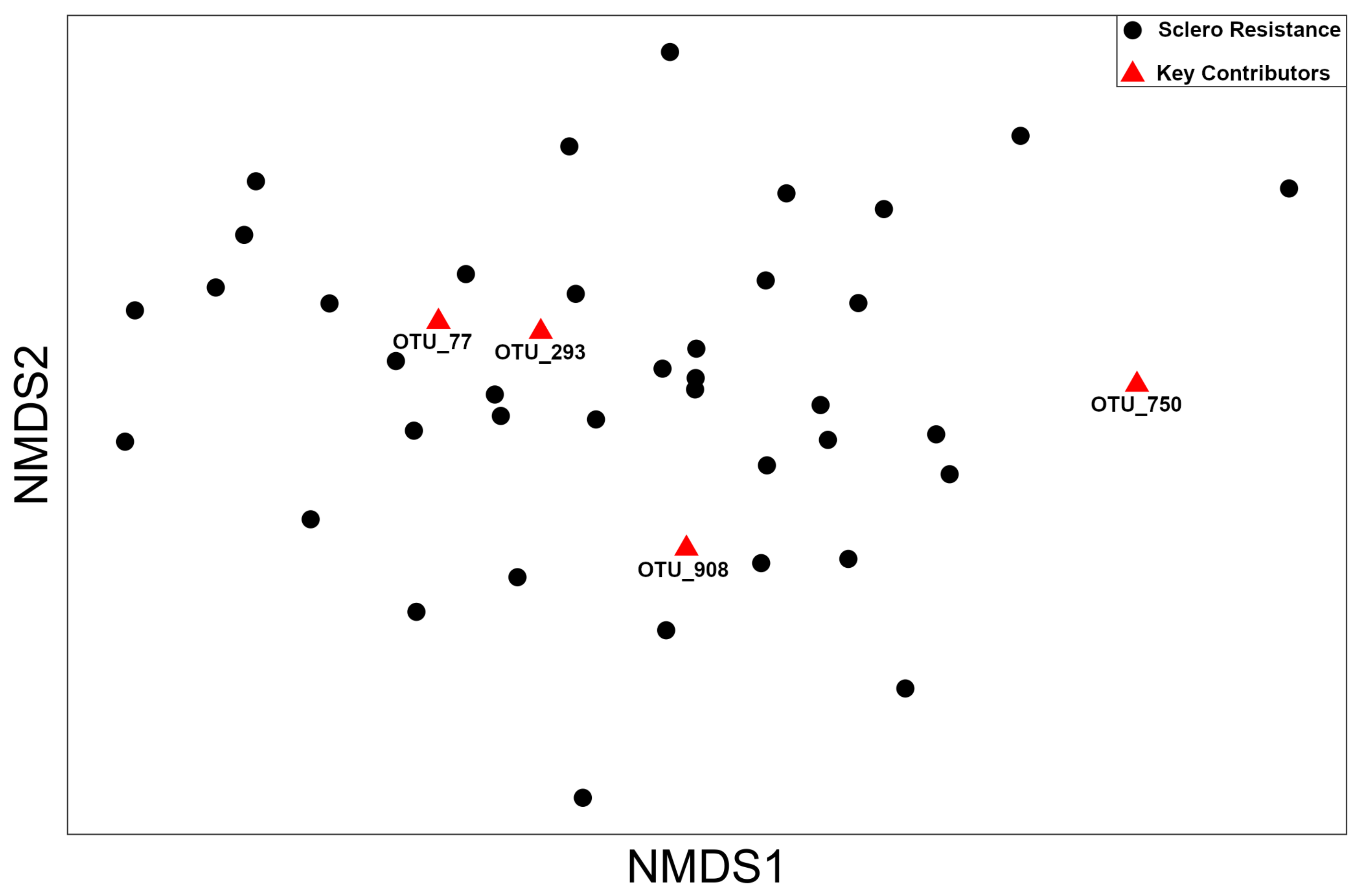

### Figure S2

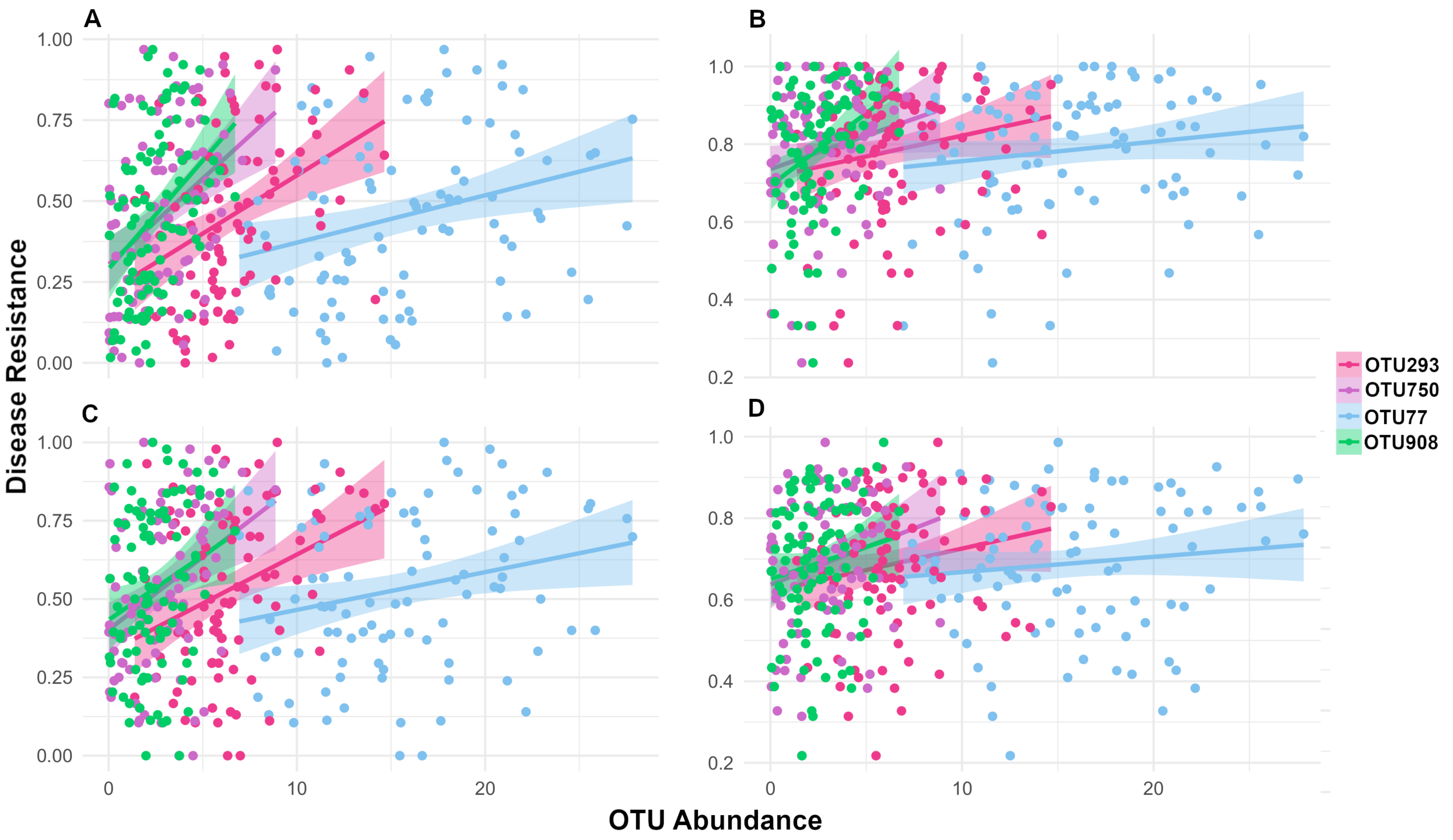

### Figure S3

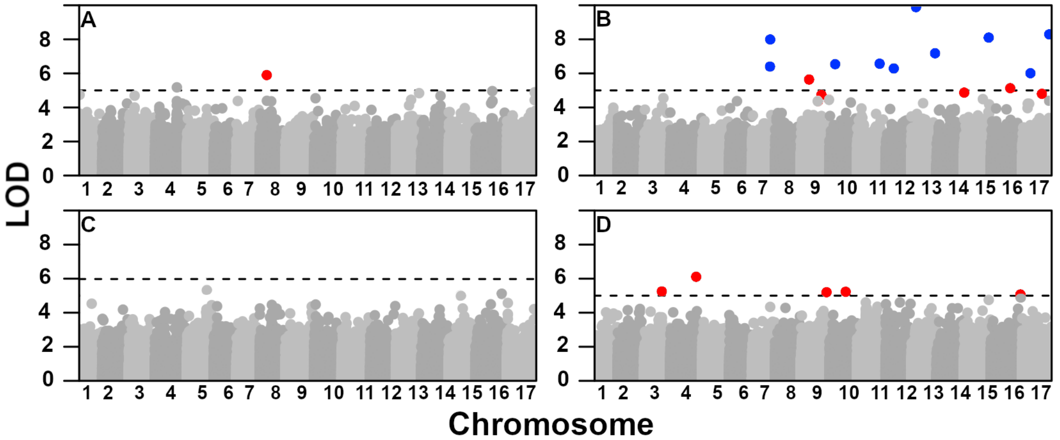

### Figure S4

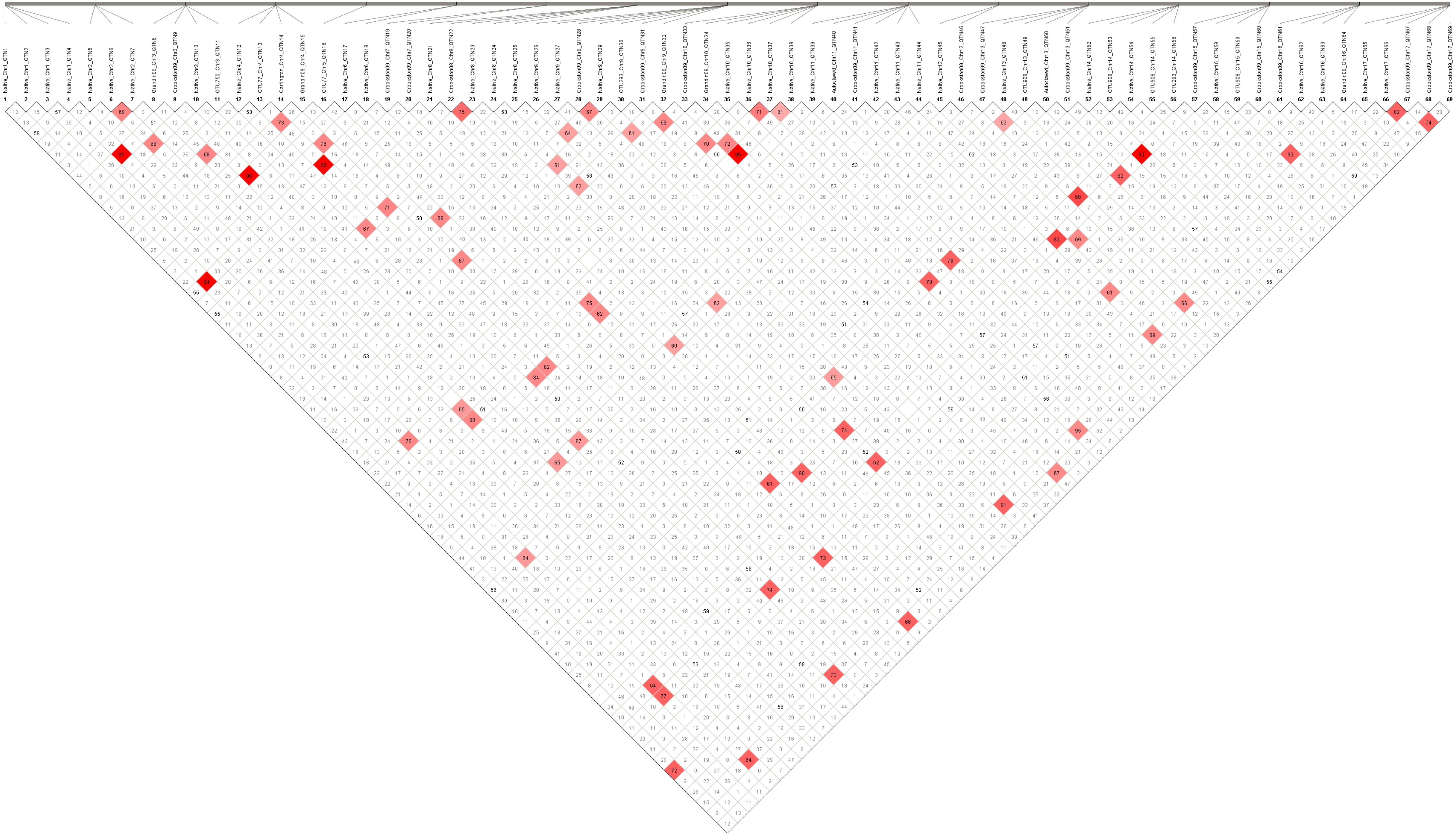
