## Supplemental Figures with captions for "Heritable associations with microbial communities are essential for necrotrophic pathogen resistance"

### Slide 1
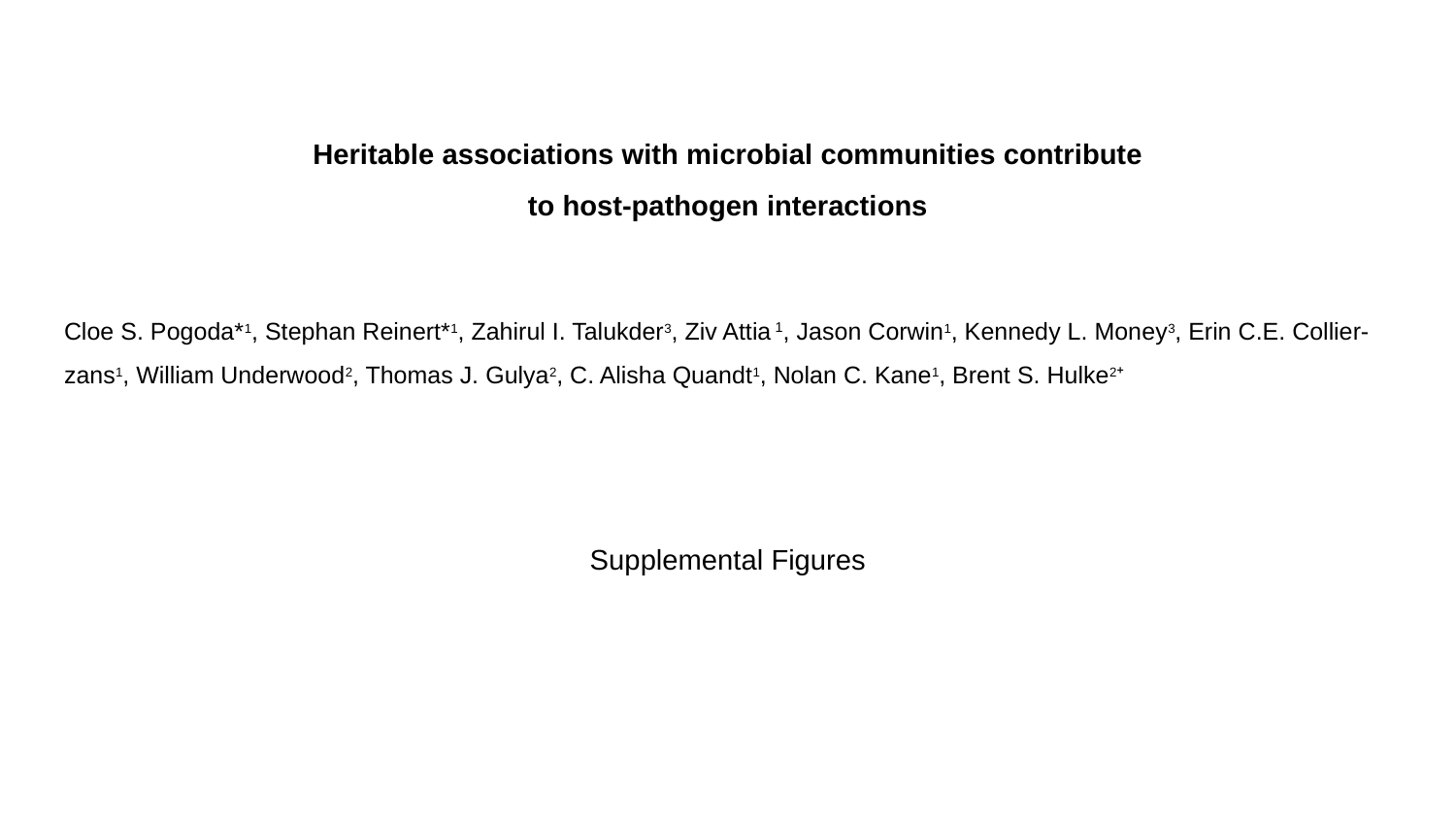

#### Heritable associations with microbial communities contribute
to host-pathogen interactions
Cloe S. Pogoda*^1^, Stephan Reinert*^1^, Zahirul I. Talukder^3^, Ziv Attia ^1^, Jason Corwin^1^, Kennedy L. Money^3^, Erin C.E. Collier-zans^1^, William Underwood^2^, Thomas J. Gulya^2^, C. Alisha Quandt^1^, Nolan C. Kane^1^, Brent S. Hulke^2^⁺
Supplemental Figures

### Slide 2
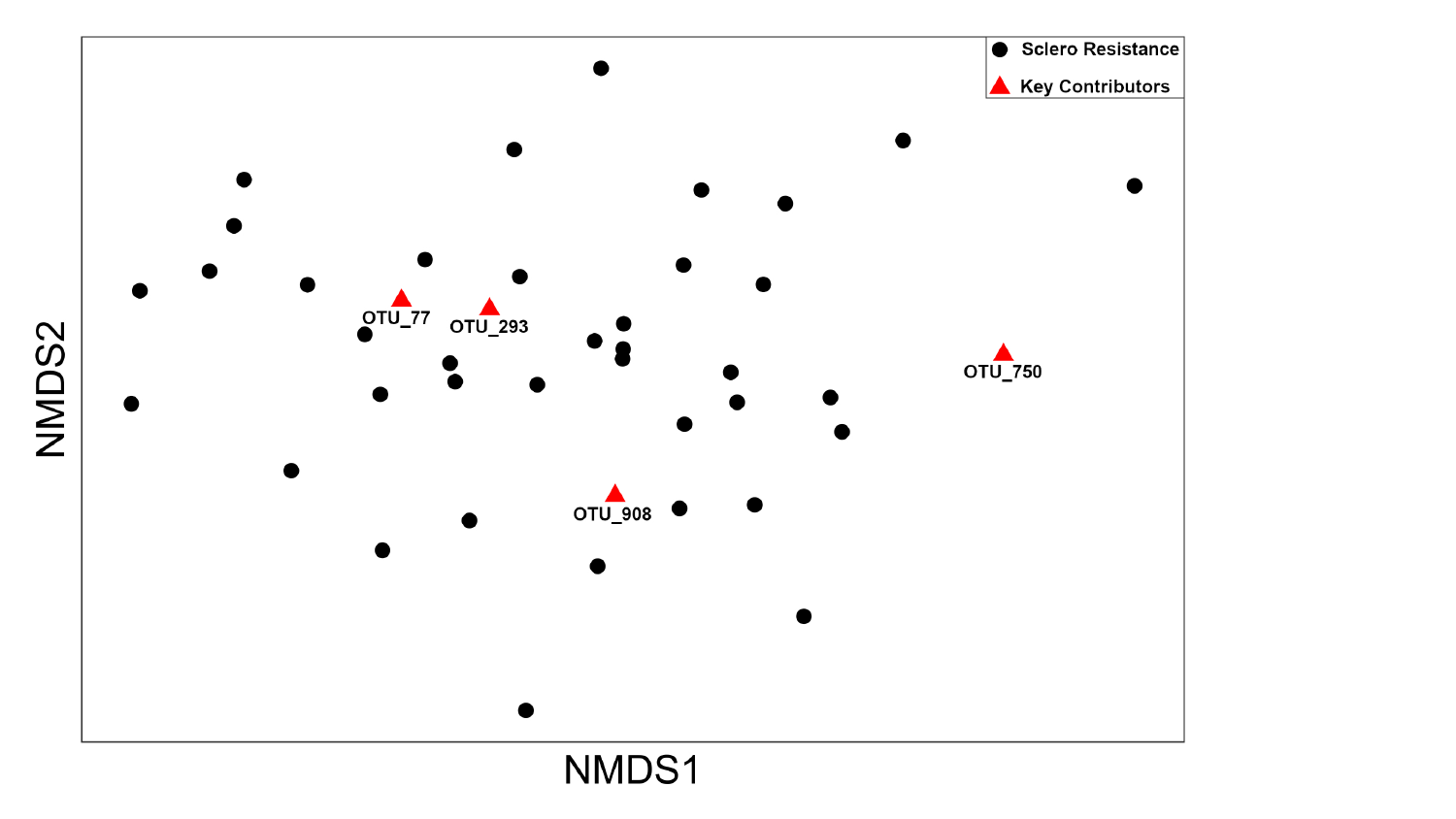

### Slide 3
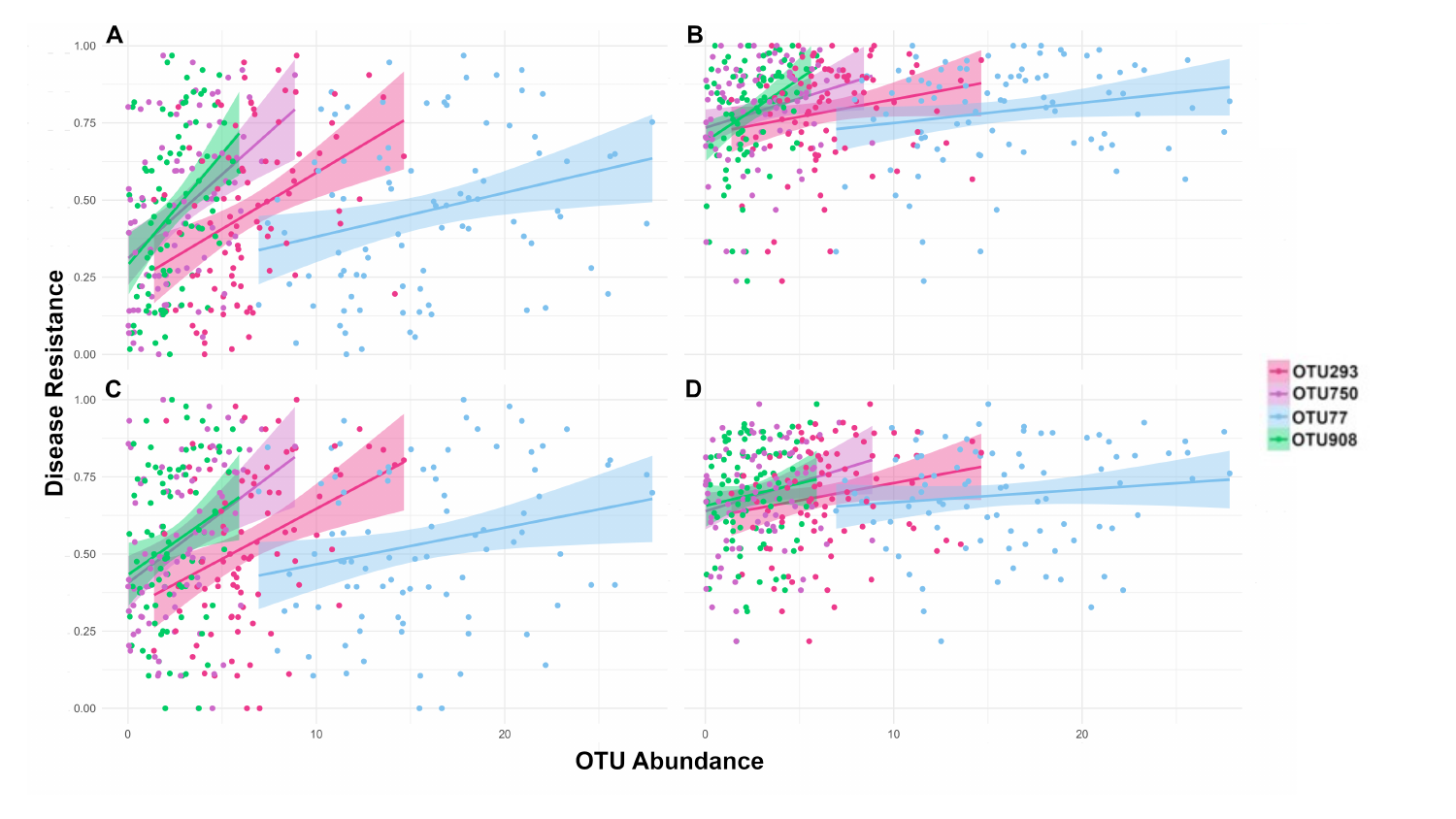

### Slide 4
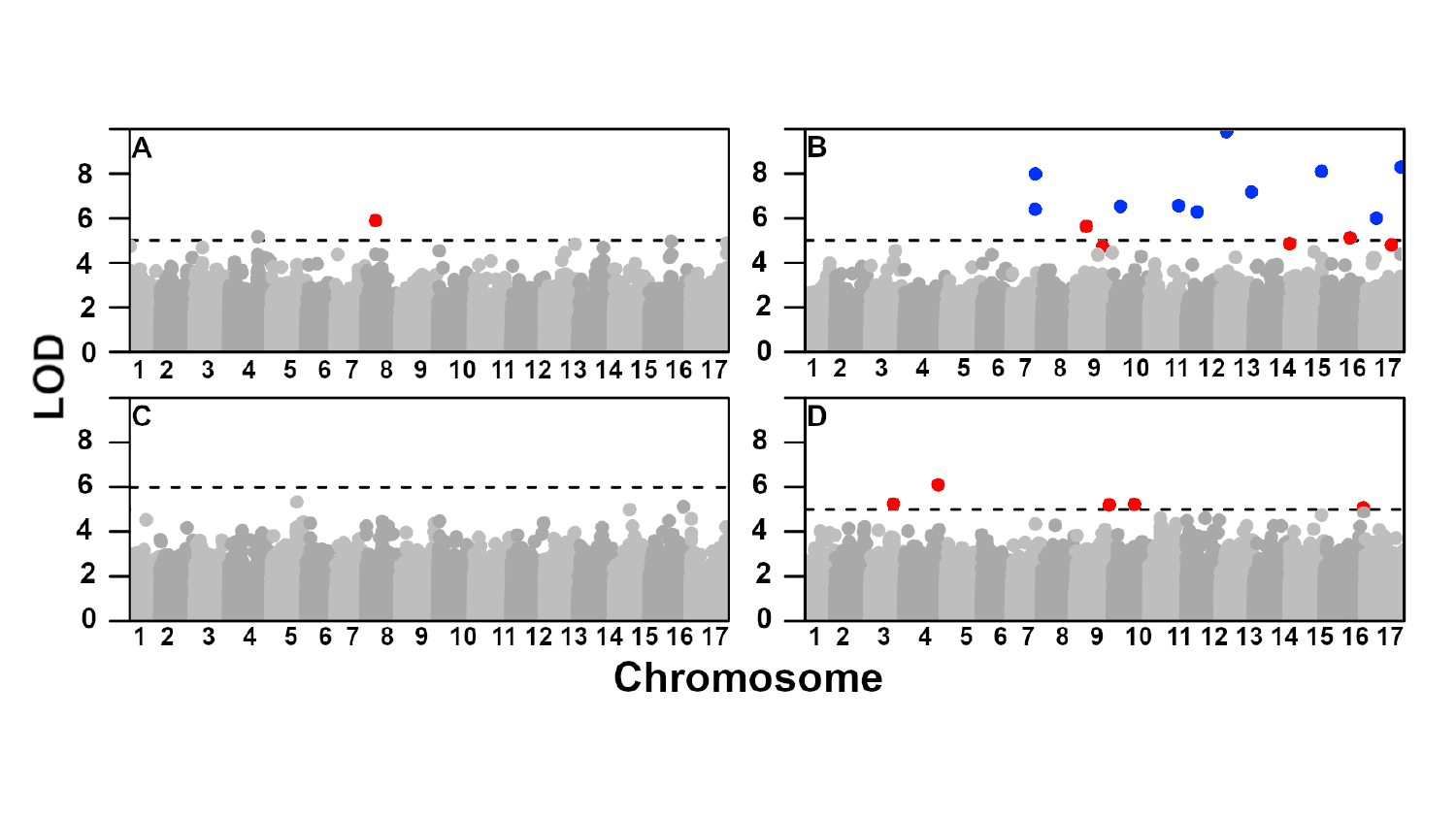

### Slide 5
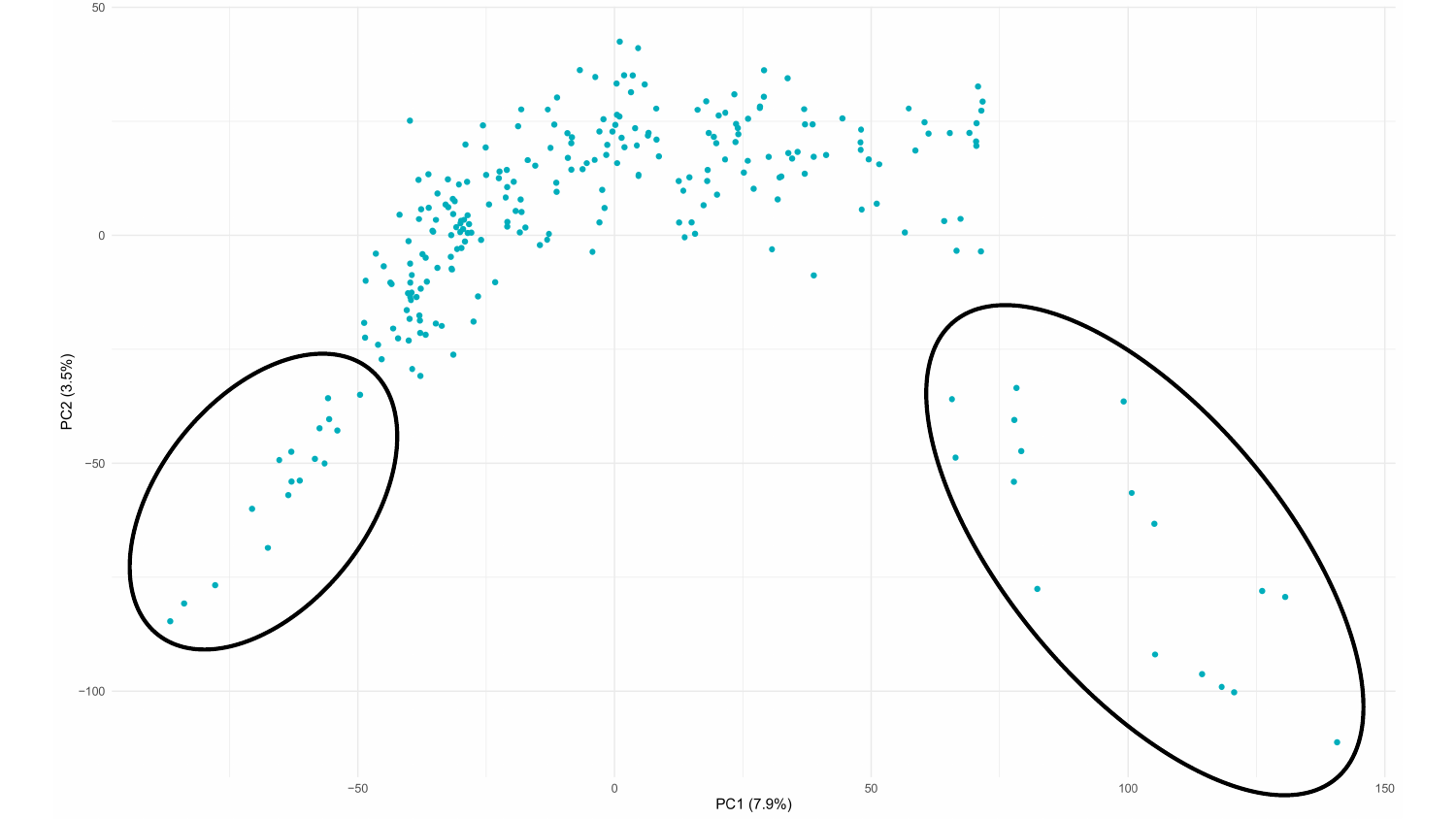

### Slide 6
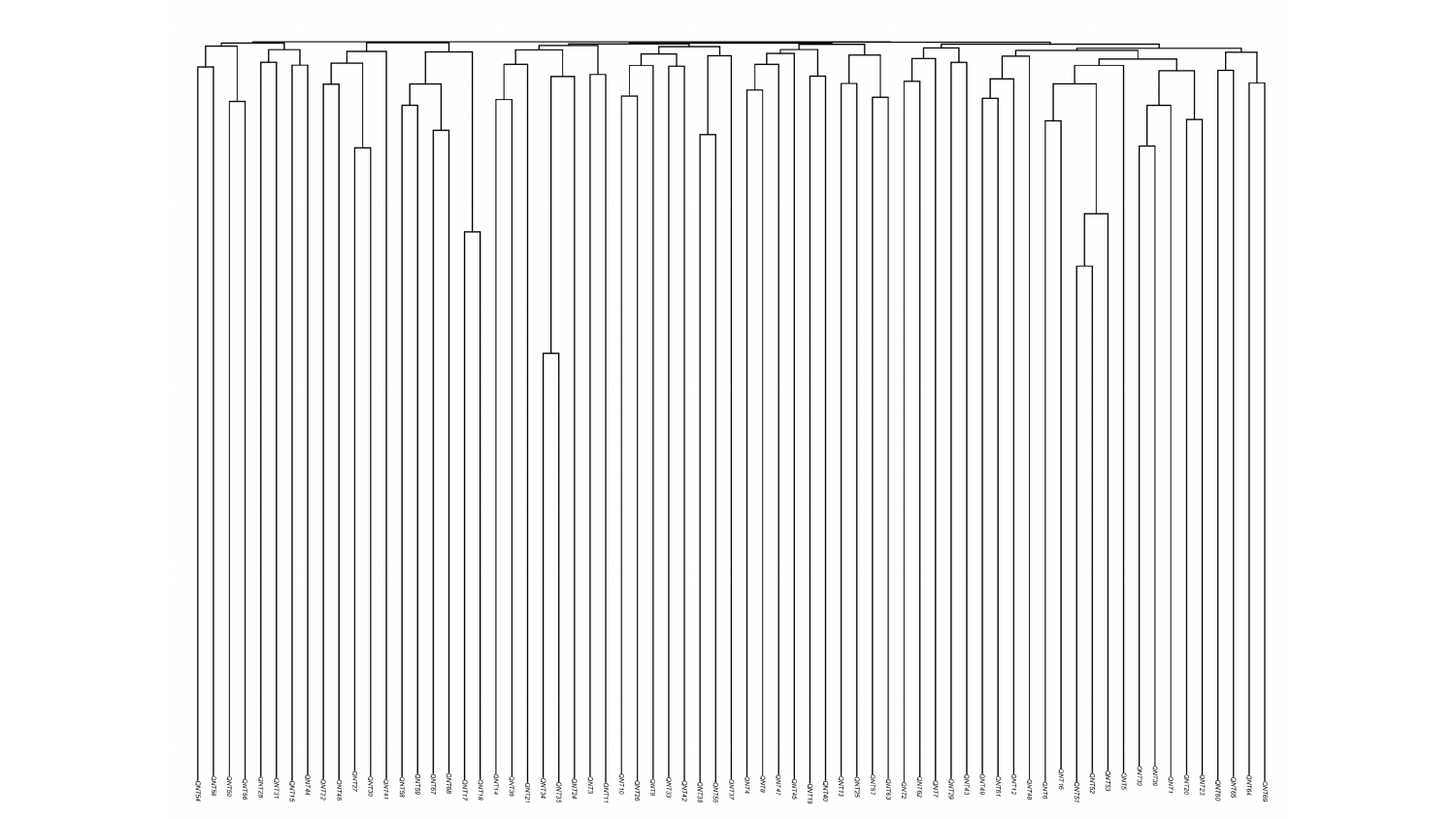
